# Multi-scale Transcriptional Characterization of HPV16 and HPV18 in Cervical Cancer: Host Pathway Alterations and Links With Viral Gene Expression Programs

**DOI:** 10.64898/2026.09.30.755713

**Authors:** Marcela Akemi Ishihara, Maiara M. Moro Valeriano, Noilson Oliveira, Izabela Mamede, Lívia Cavalcanti de Morais Gama, André Ferreira Cunha, Ícaro Maia Santos de Castro, Francisco Vásquez, Milton Yutaka Nishiyama-Jr, João Carlos Setubal, Eurico de Arruda Neto, Roger Chammas, Helder I Nakaya

## Abstract

High-risk HPV is central to cervical carcinogenesis, but whether individual genotypes drive distinct tumor biology or converge on a shared oncogenic state remains unclear. Profiling viral and host gene expression across two independent cohorts (n = 93; 76 HPV16, 17 HPV18), we show that HPV16-and HPV18-associated cervical cancers differ in host transcriptional programs. A 55-gene signature discriminated the two genotypes, and 293 genes were concordantly differentially expressed across cohorts. Pathway and transcription-factor activity inference organized these differences along genotype-specific axes: HPV16 tumors were enriched for epithelial polarity and RTK–RAS–MAPK signatures, with elevated inferred CDX2 regulon activity, a transcription factor not normally active in cervical epithelium, suggesting an aberrant lineage-identity program not previously linked to this genotype. HPV18 tumors instead showed metabolic reprogramming and inflammatory signaling (STAT3, HIF1A), alongside reduced CASP8 and increased FDX1 expression, a pattern consistent with sensitization to immunogenic cell death. Both genotypes also organized along a continuum of early-to-late viral gene expression correlating with distinct host pathways: extracellular-matrix and translation-related in HPV16, interferon/antiviral and cell-cycle/mitochondrial in HPV18, indicating that viral transcriptional state associates with host biology beyond genotype alone. These results point to candidate genotype-specific vulnerabilities that merit functional validation.

## 1 Introduction

Cervical cancer (CC) is the fourth most common malignancy among women worldwide, and persistent infection with high-risk human papillomavirus (HPV) is detected in nearly all cases, establishing HPV as the necessary etiological agent of cervical carcinogenesis^1,2^. High-risk HPV types, particularly HPV16 and HPV18, are members of the family Papillomaviridae and genus Alphapapillomavirus, belonging to the species Alphapapillomavirus 9 and Alphapapillomavirus 7, respectively. Although more than 450 HPV genotypes have been identified to date^3^, a limited number of high-risk types account for the vast majority of CC cases. Among these, HPV16 and HPV18 are responsible for approximately 77% of CCs worldwide, contributing to 61.7% and 15.3% of cases, respectively^4^. This unequal contribution suggests that the two genotypes are not equivalent in their carcinogenic potential^5,6^.

This divergence in prevalence is mirrored by differences in biological behavior and clinical presentation. HPV16 is more frequently associated with squamous cell carcinoma, whereas HPV18 has been linked to adenocarcinoma. However, this relationship is inconsistent across populations and neither genotype is restricted to a single histological subtype^7^. HPV16 infections also tend to persist longer, a major determinant of malignant progression^8^, whereas HPV18-positive tumors are associated with more aggressive clinicopathological features and poorer survival^9^.

Whether these clinical differences reflect distinct host transcriptional programs remains unresolved, in part because viral transcription is rarely analyzed alongside the host transcriptome. Although bulk RNA is capable of capturing HPV-derived transcripts^10^, these reads are commonly discarded during standard alignment to the human reference genome or used only to infer viral genotype or infection status. Previous studies have demonstrated that integrating viral and host RNA-seq data can reveal genotype-specific host transcriptional responses by jointly aligning sequencing reads to the human genome and HPV reference genomes^11^. Metatranscriptomic approaches based on taxonomic classifiers have enabled sensitive identification of viral sequences directly from unmapped RNA-seq reads^12^. These strategies provide an opportunity to simultaneously infer HPV genotype, quantify viral transcriptional activity, and investigate how viral gene expression relates to host regulatory programs.

Building on this opportunity, we developed Meta-HPV, an integrative metatranscriptomic framework that recovers and quantifies HPV transcripts directly from the unmapped fraction of bulk RNA-seq reads while simultaneously profiling host gene expression from the same dataset, requiring no additional sequencing or targeted viral capture. We validated this approach against orthogonal RT-PCR genotyping and applied it across independent HPV16-and HPV18-associated CC cohorts to characterize genotype-specific host transcriptional states, regulatory networks, and cellular remodeling processes, and to determine whether viral transcriptional activity itself, beyond genotype alone, tracks with distinct host biology. In doing so, our study establishes viral transcriptional state as an additional, informative dimension for understanding HPV-driven tumor heterogeneity, complementary to genotype and infection status.

## 2 Materials and Methods

### 2.1 Overview of the Meta-HPV pipeline

We developed Meta-HPV, a pipeline that recovers HPV transcriptional signal from the fraction of RNA-seq reads that fail to align to the human genome (typically 10 –10 reads per sample) and routinely discarded in standard analyses. Raw FASTQ reads were aligned to the GRCh38 reference genome using HISAT2 v2.2.1^13^ with default parameters, and unmapped reads were extracted with SAMtools v1.12 ^14^. These reads were classified using Kraken2 v2.1.3^15^ with default parameters, a k-mer-based taxonomic classification tool, against the Standard PlusPF database (March 2023) that includes HPV, other microbial, and human reference genome to limit false-positive assignment of human-derived reads^16^. Reads classified as HPV16 or HPV18 were realigned to the cognate reference genome (NCBI-GenBank: NC_001526.4; NC_001357.1) using HISAT2, and read counts across annotated open reading frames (E1, E2, E4, E5, E6, E7, L1, L2) were quantified with featureCounts v2.0.2^17^ to generate per-sample viral expression profiles, visualized as circular genome maps.

In parallel, mapped reads were quantified against the GENCODE v38 annotation using featureCounts (Subread v2.0) to generate host gene expression profiles, which were normalized and variance-stabilized using DESeq2 v1.34^18^. For each sample, the dominant HPV genotype was assigned based on best-matching viral read alignment, and this meta-genotype classification was used as the primary grouping variable for all downstream host and viral integrative analyses.

### 2.2 Validation and application cohorts

Meta-HPV was benchmarked against RT-PCR genotyping (targeting the L1 gene) in an independent cohort from Ure et al. (2022)^10^ comprising 446 CC RNA-seq samples (223 HPV-positive, 223 HPV-negative), including 30 RT-PCR-confirmed HPV16-positive and 14 HPV18-positive tumors. The pipeline was run blind to genotype, and classification calls were compared against RT-PCR results.

The validated pipeline was then applied to two independent, publicly available CC RNA-seq cohorts: Curty et al. (2021; n = 26; NCBI-Bioproject: PRJNA356774)^19^ and Ruiz et al. (2018; n = 67; NCBI-Bioproject: PRJNA636799)^20^, yielding a combined dataset of 93 samples (Table S1). Raw FASTQ files were retrieved from the NCBI SRA using sra-tools, quality-filtered and adapter-trimmed with fastp^21^, and processed through the Meta-HPV pipeline as above (GRCh38.p14; matching GENCODE annotation). The resulting Meta-HPV genotype classification served as the primary grouping variable for all subsequent molecular signature, meta-differential expression, regulon, and viral-program analyses.

### 2.3 Identification of a host transcriptional signature discriminating HPV16 and **HPV18**

To identify a host transcriptional signature discriminating HPV16 from HPV18 infection across both cohorts, we applied the Multi-group INTegrative (MINT) sPLS-DA framework with “regression” mode in mixOmics^22^ to TMM-normalized, log2-transformed gene counts, adjusted for study-specific effects and filtered for near-zero variance. The optimal number of components (1–5) and features per component (5–75) were selected by tuning on balanced error rate (BER) and centroid-distance classification with three repeated cross-validations. Model performance was assessed by AUROC of the first discriminant component (perf function); the signature was defined as the union of features selected across components (selectVar), ranked by VIP score. Sample separation was visualized by UMAP on the MINT sPLS-DA latent scores. For visualization purposes, log2 fold-changes (log2FC) for the 55 signature genes were calculated separately with edgeR (v4.8.2) (v4.8.2)^23^, independent of the meta-analysis differential expression results described in Section 2.4.

### 2.4 Meta-analysis of differential gene expression across cohorts

Differential expression between HPV16-and HPV18-positive samples was performed independently in each cohort using DESeq2 (v1.34)^18^, with sequencing batch as a covariate where applicable. Study-level results were integrated using a random-effects meta-analysis model in MetaVolcanoR (Marinho et al., submitted; https://github.com/csbl-usp/MetaVolcanoR), which accounts for between-study heterogeneity. Genes with |summary log2FC| > 1 and adjusted meta-analysis p < 0.05 were considered consistently differentially expressed and visualized as a meta-volcano plot, with sign-consistency scores (±2) indicating full directional concordance across studies.

### 2.5 Transcription factor regulon and pathway activity analysis

Genes were ranked by meta-analysis effect size (randomSummary), with positive values reflecting higher expression in HPV16. TF activity was inferred from DoRothEA regulons (confidence levels A-C)^24^ using the univariate linear model (ULM) method in decoupleR^25^, with positive/negative ULM scores indicating higher regulon activity in HPV16/HPV18, respectively, and significance assessed by Benjamini–Hochberg-corrected FDR < 0.05. The same ranked gene list and a Multivariate Linear Model (MLM) framework were applied to MSigDB Hallmark gene sets^26^ to infer pathway-level activity, with positive/negative scores indicating enrichment in HPV16/HPV18 tumors.

### 2.6 Viral transcriptional programs and associated host pathways

Samples were aligned to HPV16 and HPV18 genomes using HISAT2^27^. For each FASTQ file, reads were aligned in unspliced mode using the HPV16 reference index, with up to 10 reported alignments per read (-k 10). Alignment sensitivity was controlled using the scoring function --score-min L,0,-0.6, and spliced alignment was disabled (--no-spliced-alignment). Gene-level read counts were subsequently obtained using featureCounts^17^. For each genotype (HPV16 and HPV18), an early score (E2 + E5) and a late score (L1 + L2) were computed per sample from log1p-transformed CPM values. E2 and E5 were selected as representative early genes because E6 and E7 showed uniformly high expression across nearly all samples regardless of transcriptional stage, limiting their utility for distinguishing early-from late-dominant profiles, whereas E1 and E4 were excluded due to multi-mapping ambiguity during alignment (data not shown). E2 and E5 were therefore retained as the most conservative markers of early gene expression. A residual score was calculated as the difference between the late and early scores, with positive values indicating a late-dominant bias and negative values indicating an early-dominant bias. Visual inspection of this score suggested three loosely bounded profiles (early-dominant, mixed, and late-dominant) although no formal clustering procedure was applied, and the residual score was treated as continuous in all downstream analyses. Host genes associated with this early-to-late continuum were identified using a quasi-likelihood edgeR^23^ model (residual score as a continuous covariate, study as a blocking factor) and by Spearman correlation with normalized host gene expression. Genes were ranked by signed −log10(p-value) from the Spearman correlation and tested for pathway enrichment using fgseaMultilevel against the Reactome database^28^, with significant pathways defined as FDR < 0.05 and fewer than 100 member genes.

## 3 Results

### 3.1 Meta-HPV enables HPV Genotype identification from standard RNA-seq data

Meta-HPV was applied to an independent validation cohort of 446 CC RNA-seq samples (223 HPV-positive, 223 HPV-negative), including 30 RT-PCR-confirmed HPV16-positive and 14 HPV18-positive tumors^29^. Meta-HPV recovered HPV-derived transcripts matching the RT-PCR-assigned genotype in every case: HPV16-positive tumors showed high HPV16-aligned read counts with minimal HPV18 signal, and HPV18-positive tumors showed the reciprocal pattern (Figure 1B), confirming the pipeline’s ability to accurately genotype HPV directly from bulk RNA-seq data.

**Figure 1.**
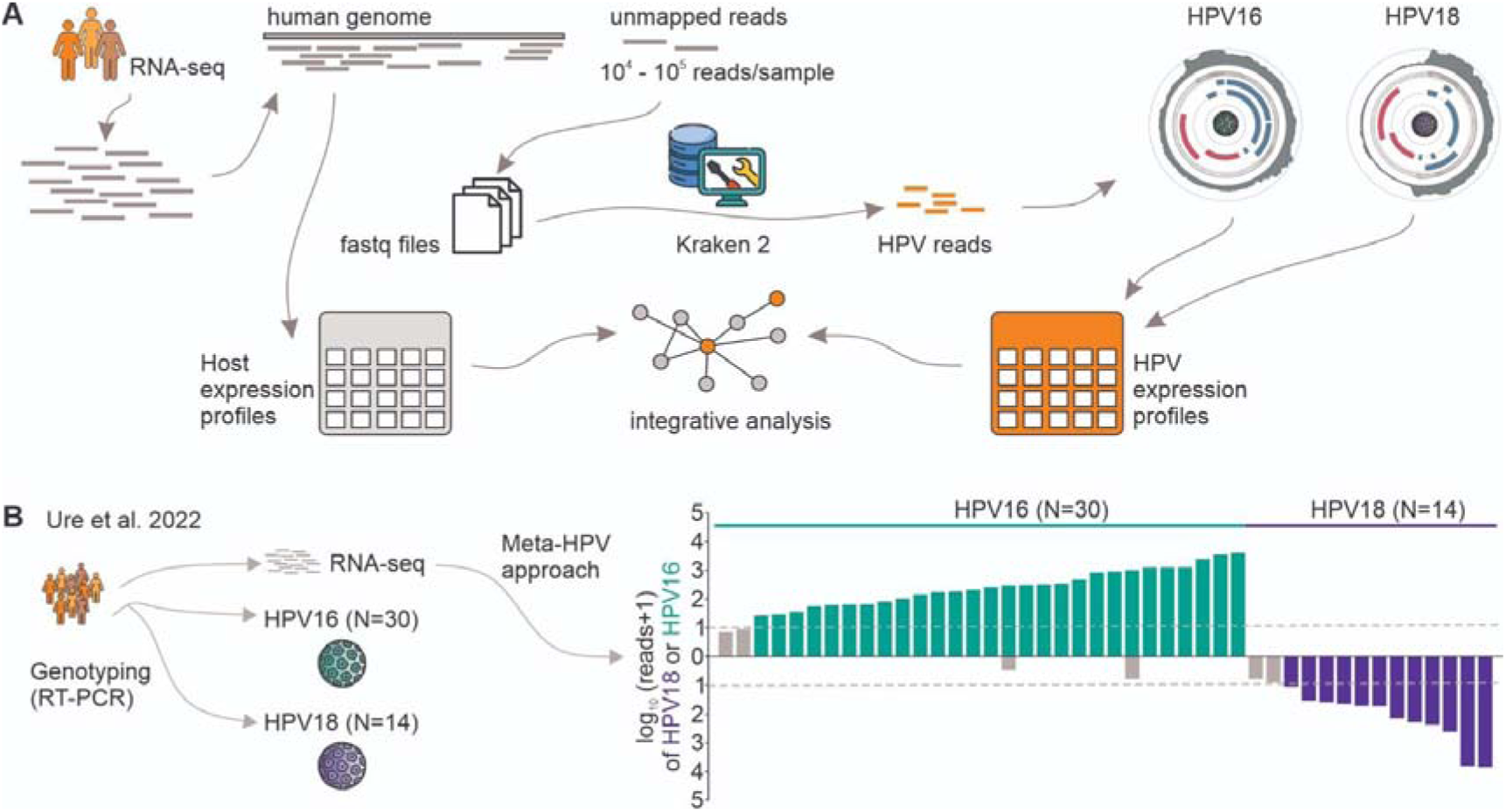
Overview of the Meta-HPV approach and validation in an independent cohort. (A) Schematic of the Meta-HPV pipeline. RNA-seq reads are aligned to the human reference genome, and unmapped reads (10 –10 reads per sample) are extracted and classified using Kraken 2 to identify HPV-derived sequences. Classified HPV reads are used to construct HPV expression profiles, which are integrated with host expression profiles derived from the mapped reads for downstream integrative analysis. Circular genome maps illustrate the resulting HPV gene expression profiles for HPV16 and HPV18. (B) Validation of the Meta-HPV approach in the Ure et al. (2022) cohort, comprising 30 HPV16-and 14 HPV18-positive samples genotyped by RT-PCR. The bidirectional bar plot shows log-transformed read counts (log (reads + 1)) for HPV18 (above axis, teal) and HPV16 (below axis, purple) per sample, demonstrating concordance between Meta-HPV classifications and orthogonal RT-PCR genotyping.

Having established the ability of Meta-HPV to reliably assign HPV genotypes from RNA-seq data, we next investigated whether HPV16-and HPV18-positive tumors exhibit distinct host transcriptional landscapes, applying MINT sPLS-DA across two independent, publicly available cohorts (Curty et al., 2021; Ruiz et al., 2018; combined n = 93: 76 HPV16-positive, 17 HPV18-positive tumors). Applying MINT sPLS-DA to the combined cohort (76 HPV16-positive, 17 HPV18-positive tumors) yielded a 55-gene host transcriptional signature (50 features on the first component, five on the second; Figure 2B) that discriminated HPV16 from HPV18 tumors, with a cross-validated AUC of 0.77 (leave-one-out; p = 4.5 × 10). UMAP projection of the latent components showed separation by genotype rather than by study of origin (Figure 2C), and the signature was consistently recovered

**Figure 2.**
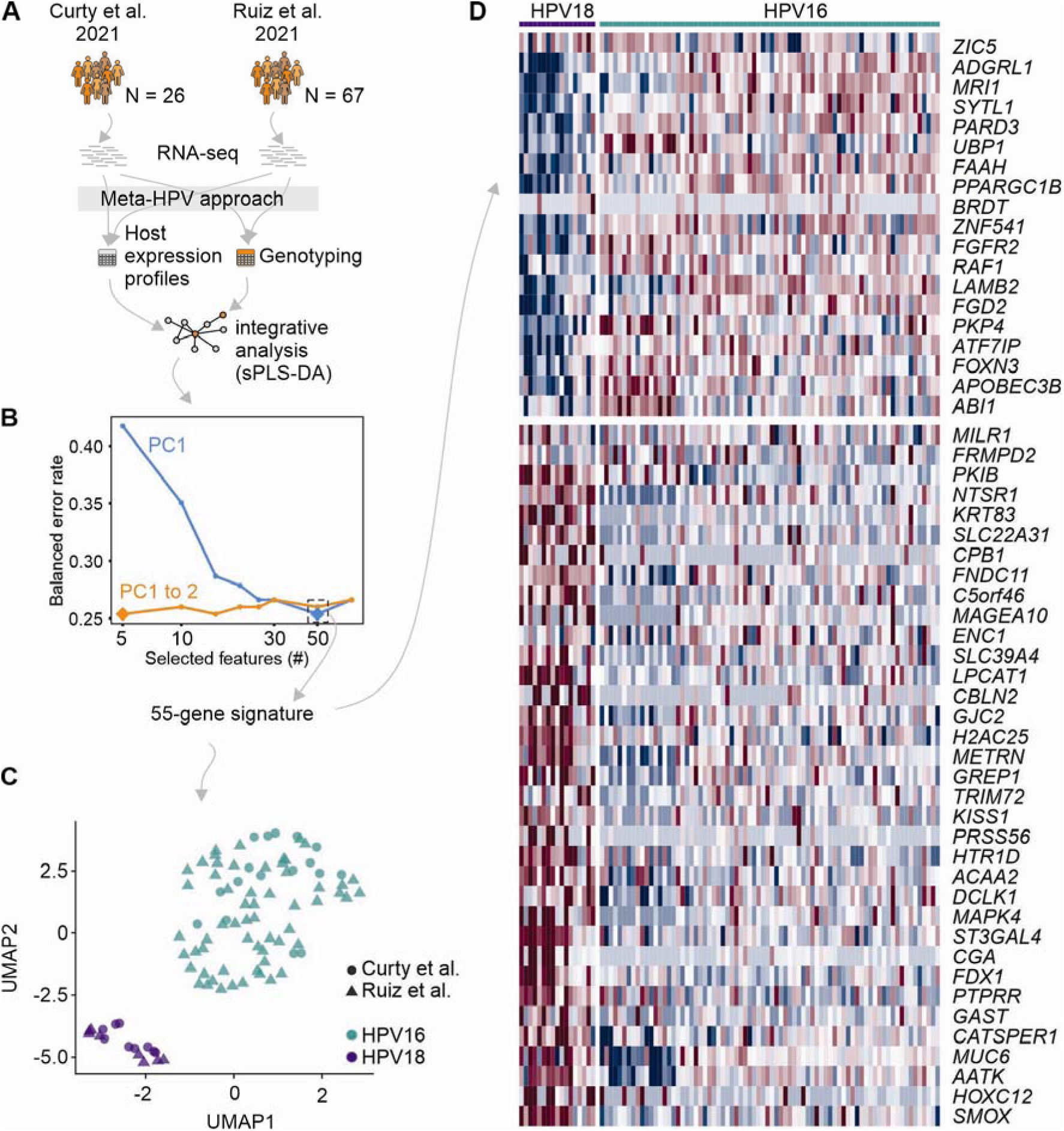
Integrative transcriptomic analysis identifies an HPV genotype–associated gene signature across independent bulk RNA-seq cohorts. (A) Schematic overview of the Meta-HPV workflow. Bulk RNA-seq data from two independent cohorts (Curty et al., 2021, n = 26; Ruiz et al., 2018, n = 67) were processed using the Meta-HPV approach to simultaneously derive host (human) gene expression profiles and determine HPV genotype. Samples classified as HPV16 or HPV18 were used for integrative multi-study analysis with MINT sparse partial least squares discriminant analysis (MINT sPLS-DA), using HPV genotype as the categorical outcome variable. (B) Model tuning results showing the balanced error rate (BER) as a function of the number of selected features. Performance was evaluated across latent components, and optimization of the number of components and selected features identified a 55-gene signature with the lowest classification error. (C) UMAP projection of sample latent scores derived from the optimized MINT sPLS-DA model using the 55-gene signature. Samples cluster according to HPV genotype (HPV16 versus HPV18) across both cohorts, indicating robust genotype-associated transcriptional differences independent of study origin. (D) Heatmap of the 55-gene signature across all samples. Gene expression values are represented as row-wise normalized log-fold change (logFC) values, highlighting distinct expression patterns between HPV16-and HPV18-positive tumors. Columns represent individual samples grouped by genotype, and rows represent signature genes selected by the MINT sPLS-DA model.

The 55-gene signature showed coordinated, genotype-specific expression patterns (Figure 2D). Genes with higher expression in HPV18 tumors included ZIC5, ADGRL1, and FGFR2 (epithelial regulation and signaling), while genes most associated with HPV16 included KRT83, SLC39A4, and TRIM72 (epithelial differentiation and cellular organization).

### 3.2 HPV16 and HPV18 tumors exhibit distinct, coordinated transcriptional **programs**

Meta-differential expression analysis across both cohorts identified 376 genes significantly differentially expressed (DE) between HPV16-and HPV18-positive samples (randomSummary > |1|; FDR < 0.05). Among these, 293 genes (78%) exhibited a sign consistency score of ±2, indicating complete concordance in the direction of differential expression across both datasets (Figure 3A; Table S2). Of the concordantly DE genes, 123 were upregulated and 170 were downregulated, with effect sizes ranging from −5.07 to 5.45 (median randomSummary = −1.12). The most positively associated genes included ENSG00000288047 (randomSummary = 5.45, FDR = 0.024), CLEC2A (4.55, FDR = 0.0077), ADH1C (4.46, FDR = 2.96 × 10⁻⁵), HMX1 (4.31, FDR = 0.045), and ENSG00000287594 (4.26, FDR = 0.026). Additional highly ranked genes with positive effect sizes were ZPBP2, CRNN, SCGB2A2, PIGR, and PNLIPRP3. Conversely, the strongest negative associations were observed for KRT83 (randomSummary = −5.07, FDR = 0.0061), SLC6A3 (−4.48, FDR = 0.0012), CT83 (−4.42, FDR = 0.033), ENSG00000234476 (−4.19, FDR = 0.025), and NTM-AS3 (−4.19, FDR = 0.032), followed by GREP1, SFTA1P, MAGEA10, CPB1, and C5orf46.

**Figure 3.**
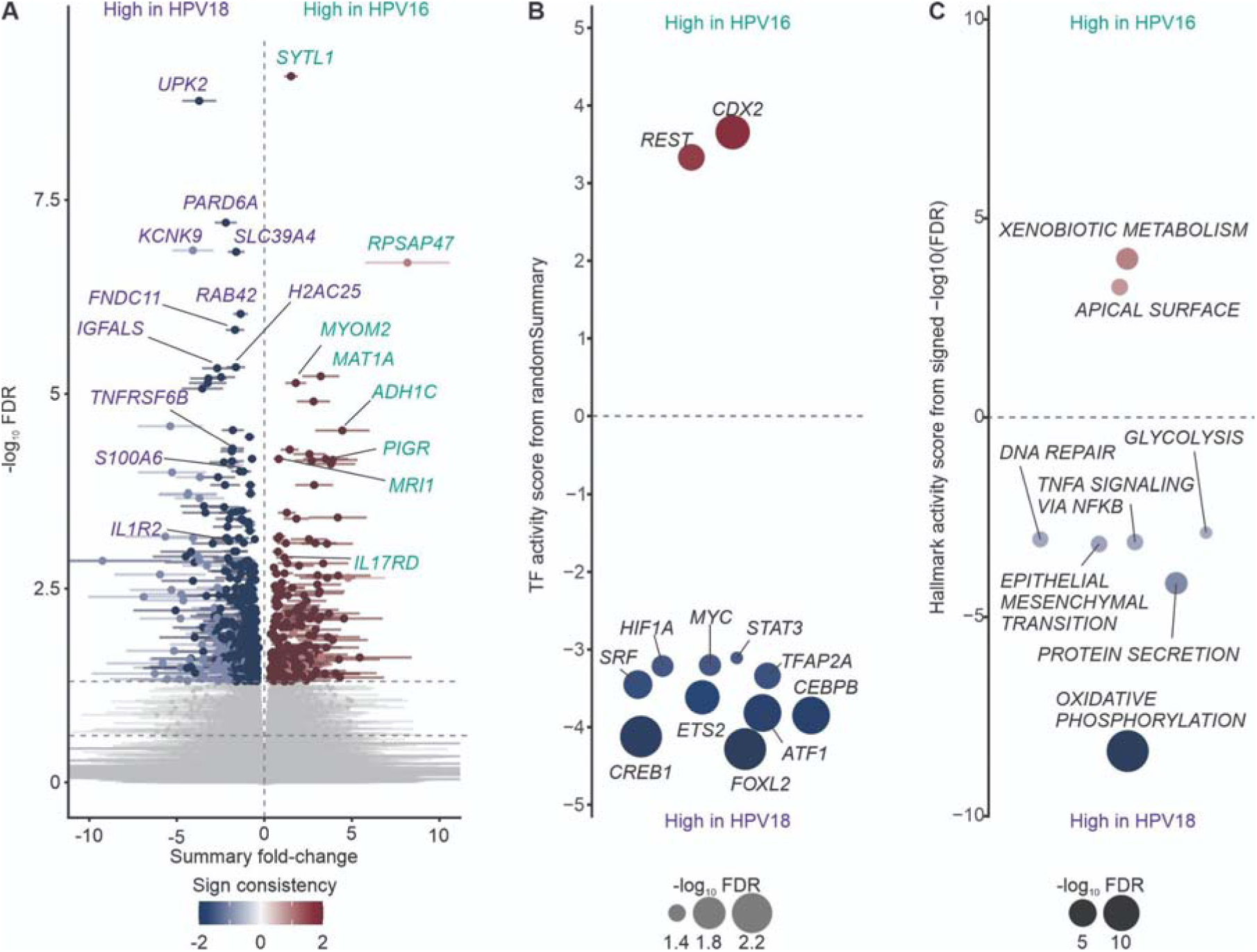
Multi-layer meta-analysis reveals consistent transcriptional, regulatory, and pathway-level differences between HPV16-and HPV18-positive tumors. (A) Meta-volcano analysis identifying genes that were consistently differentially expressed between HPV16-and HPV18-positive tumors across both cohorts. Differential expression was performed independently in each study and integrated to identify genes exhibiting concordant directionality and significance. Only features passing an adjusted P value threshold of < 0.05 were retained. The color scale indicates sign consistency across studies, where −2 represents genes significantly downregulated in both studies and +2 represents genes significantly upregulated in both studies. Representative top-ranked genes are labeled. (B) Differential transcription factor (TF) activity analysis inferred from gene expression profiles and integrated across studies. TF regulons with adjusted P values < 0.05 are shown. Positive activity scores indicate increased regulon activity in HPV16-positive tumors, whereas negative scores indicate increased activity in HPV18-positive tumors. Multiple TF regulons displayed significant genotype-associated differences, with a larger proportion exhibiting HPV18-biased activity. (C) Cancer hallmark enrichment analysis comparing pathway activity between HPV16-and HPV18-positive tumors. Hallmarks were evaluated across studies and integrated using the same meta-analytic framework, retaining only significant features (adjusted P < 0.05). Positive enrichment scores indicate pathways preferentially associated with HPV16-positive tumors, whereas negative enrichment scores indicate pathways preferentially associated with HPV18-positive tumors. Consistent with the transcription factor analysis, a greater number of cancer hallmarks exhibited HPV18-biased activity, highlighting broader pathway-level differences between HPV genotypes.

Several of the top-ranking genes with positive effect sizes, including CLEC2A, PIGR, CRNN, and ADH1C, are associated with epithelial differentiation, mucosal immune responses, and metabolic regulation. In contrast, genes with strong negative effect sizes, such as KRT83, SLC6A3, and MAGEA10, may reflect distinct transcriptional programs that distinguish HPV16 from HPV18 infection and underscores substantial differences in host cellular responses between the two viral types.

TF activity inference (DoRothEA regulons, ULM/decoupleR) identified TFs with strong differential activity between genotypes (Figure 3B), with FDR < 0.05. CDX2 and REST showed the highest HPV16-biased activity (scores = 3.67 and 3.31), both canonically linked to differentiation and growth control. A larger number of TFs, including FOXL2, CREB1, CEBPB, ATF1, ETS2, SRF, TFAP2A, HIF1A, MYC, and STAT3, showed HPV18-biased activity, with FOXL2 (−4.28), CREB1 (−4.12), and CEBPB (−3.85) the strongest (Table S3). Several of these (CREB1, CEBPB, MYC, STAT3, HIF1A) are established regulators of proliferation, metabolism, and stress signaling.

Hallmark pathway activity analysis (MSigDB, ULM/decoupleR) identified several pathways with significant differential activity (Figure 3C). Xenobiotic Metabolism and Apical Surface showed the strongest HPV16-biased enrichment (scores = 3.99 and 3.27), reflecting detoxification and epithelial surface organization programs. A larger number of pathways were HPV18-biased, led by Oxidative Phosphorylation (score = −8.38, FDR = 2.85 × 10⁻¹⁵), followed by Protein Secretion (−4.15), Epithelial–Mesenchymal Transition (−3.16), and TNFα Signaling via NFκB (−3.13) (Table S4), consistent with substantial metabolic and inflammatory reprogramming in HPV18-associated tumors.

### 3.3 Viral Metatranscriptomic Profiling Reveals Distinct HPV Gene Expression Programs Associated with Specific Host Biological States

Consistent with their central role in oncogenic transformation, E6 and E7 were the most highly expressed HPV genes in both genotypes (Figure S1), motivating their exclusion from the early/late transcriptional score described below (see Methods). Circular genome maps of normalized viral transcript abundance revealed subtype-specific expression landscapes, with HPV16 and HPV18 displaying distinct patterns of early and late gene activity across their genomes (Figure 4A,B). Projecting samples onto an early-to-late transcriptional axis, defined by the relative expression of the early regulatory genes E2 and E5 and the late structural genes L1 and L2, revealed three loosely bounded transcriptional profiles along this continuum in both HPV16 and HPV18 cohorts, rather than sharp, well-separated clusters (Figure 4B): an early-dominant profile, a late-dominant profile, and an intermediate mixed profile with concurrent expression of both programs, centered near a residual score of zero. This organization was consistent across both HPV subtypes.

**Figure 4.**
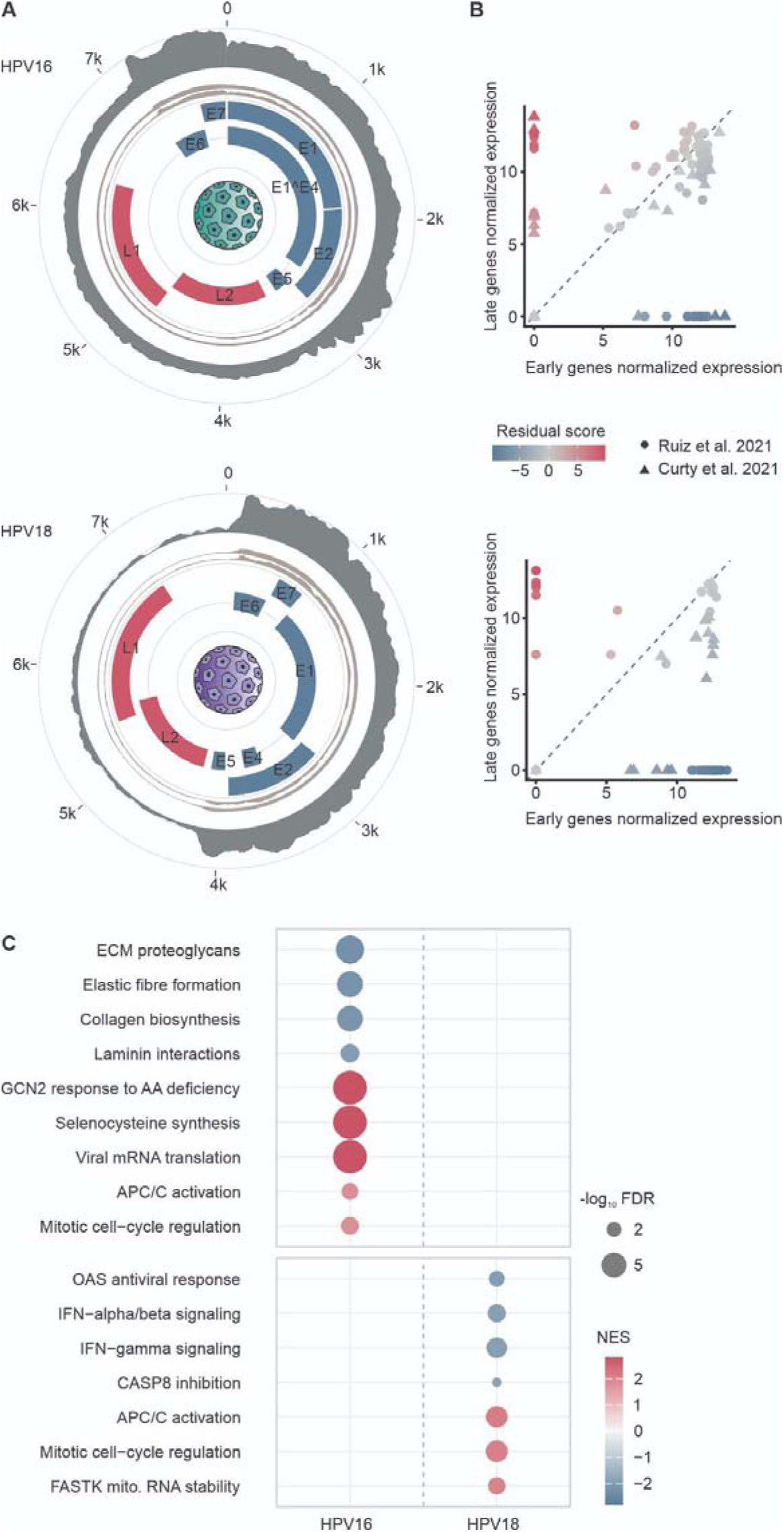
Distinct HPV16 and HPV18 viral transcriptional programs are associated with genotype-specific host biological processes. (A) Circular plots of viral gene expression from metatranscriptomic RNA-seq data in HPV16-and HPV18-positive tumors. Outer histograms show combined viral transcript abundance across both cohorts; inner tracks show study-specific profiles (Curty et al. and Ruiz et al.). Early genes (E1–E7) are shown in blue, late genes (L1–L2) in pink. (B) Classification of samples by viral transcriptional state, based on normalized expression of early (E2, E5) and late (L1, L2) viral genes, for HPV16-positive (top) and HPV18-positive (bottom) tumors. Each sample was assigned a residual score reflecting deviation from the expected early–late relationship, with positive values indicating a late-dominant program and negative values indicating an early-dominant program. Point color denotes residual score; symbols denote study origin. (C) fgsea of Reactome pathways associated with the viral program residual score, shown separately for HPV16 and HPV18. Dot size reflects enrichment significance (−log10 FDR). Color indicates normalized enrichment score (NES).

Despite this shared organizational structure, the host biological processes associated with each viral transcriptional state differed markedly between HPV16 and HPV18. In HPV16-infected samples, the early gene-dominant state was enriched for extracellular matrix (ECM)-related pathways, including ECM proteoglycans, elastic fibre formation, collagen biosynthesis, and laminin interactions (Figure 4C). The late gene-dominant state was instead associated with pathways related to host translational capacity and biosynthetic activity, including viral mRNA translation, selenocysteine synthesis, the GCN2 response to amino acid deficiency, APC/C activation, and mitotic cell-cycle regulation (Figure 4C).

A distinct pattern was observed in HPV18-infected samples. The early gene-dominant state showed enrichment of antiviral and immune-response pathways, including interferon-α/β signaling and interferon-γ signaling, OAS-mediated antiviral responses, and CASP8-associated apoptotic regulation. The late gene-dominant state was enriched for pathways related to cell-cycle progression and mitochondrial function, including APC/C activation, mitotic cell-cycle regulation, and FASTK-mediated mitochondrial RNA stability (Figure 4C) (Table S5).

## 4 -Discussion

Despite the central role of high-risk HPV in cervical carcinogenesis, it has remained unclear whether individual genotypes drive distinct tumor transcriptional programs or converge on a shared oncogenic state. By profiling viral and host transcription together across independent cohorts, we show that HPV16-and HPV18-associated CCs occupy divergent biological states within a single histological context.

This was made possible by Meta-HPV, which recovers HPV transcripts from RNA-seq reads that fail to align to the human reference and achieved perfect concordance with orthogonal RT-PCR genotyping (Figure 1B), with the resulting host classifier reaching a cross-validated AUC of 0.77. Because the recovered signal is transcriptional, it reports active viral gene expression rather than past exposure^11^, consistent with prior work showing metatranscriptomics can detect HPV in samples called negative by PCR^30^. Including the human genome alongside viral and microbial references in the classification database further guards against a documented failure mode of k-mer–based classifiers, which can misassign human sequence to microbial taxa when the reference database is incomplete and thereby inflate spurious microbiome–cancer associations^16^. Together, these design choices support that the genotype-and state-specific findings below reflect genuine biology rather than assay artifact.

HPV16 tumors were marked by an epithelial and stem-like identity program (ECM remodeling, apical-surface organization, AhR-linked xenobiotic activity, and CDX2/REST regulon activity) consistent with the biology of squamous cell carcinoma^31^ (Figure 3B). HPV16 E6 and E7 oncoproteins can induce an aberrant cadherin switch, downregulating E-cadherin and upregulating N-cadherin to promote a migratory, invasive phenotype^32^, and the same oncoproteins target the basolateral polarity regulators Dlg and Scribble, producing aberrant spindle orientation and tissue multilayering^33^. This is reinforced at the signature level: PARD3, a core determinant of apical-basal polarity and an established tumor suppressor in other squamous malignancies^34^, was independently recovered among the HPV16-associated classifier genes, as were FGFR2 and RAF1 (CRAF) (Figure 2D), which implicate the druggable RTK–RAS–MAPK axis as a second, genotype-specific vulnerability with existing clinical-stage inhibitors^35^. APOBEC3B, also among the discriminative classifier features, points to a further, complementary route to genomic instability in this genotype, consistent with prior reports of APOBEC3B mutational signatures in HPV16 but not HPV18 lesions^36^. At the regulatory level, CDX2 showed the strongest HPV16-biased transcription factor activity of any TF tested. CDX2 is the master determinant of intestinal-type epithelial identity and is not physiologically expressed in normal cervical epithelium. Prior immunohistochemical work has documented aberrant CDX2 expression in a subset of cervical adenocarcinomas, particularly the endometrioid and endocervical usual-type subtypes, even in the absence of overt colorectal morphology^37^. Its elevated regulon activity here, in a genotype driving predominantly squamous rather than glandular disease, suggests a degree of aberrant lineage-identity signaling that has not been classically associated with HPV16-driven carcinogenesis and warrants dedicated functional follow-up. HPV18 tumors, in contrast, showed broad metabolic reprogramming (oxidative phosphorylation, glycolysis), epithelial-mesenchymal transition, and an inflammatory circuit coordinated by STAT3, HIF1A, MYC, CREB1, and CEBPB^38,39^, consistent with functional evidence that HPV18 genomic integration itself can drive glycolytic reprogramming and NF-κB/MAPK activation^40^. Two classifier genes point to a mechanistically coupled, targetable vulnerability layered on top of this circuit. CASP8, which normally suppresses RIPK1/RIPK3/MLKL-dependent necroptosis in addition to its apoptotic role, was inhibited specifically during the early viral gene-dominant state. Low CASP8 is itself linked to poor CC prognosis^41^. This may sensitize HPV18 tumors to a highly inflammatory, immunogenic form of cell death rather than simply preventing immune clearance. This co-occurred with enrichment of FDX1, a master regulator of cuproptosis whose induction is itself potentiated by interferon-γ signaling via the STAT1–IRF1 axis^42^, and whose associated cell death is itself immunogenic, together nominating copper ionophores combined with checkpoint blockade as a genotype-informed combination strategy worth testing in HPV18-positive disease, in line with a broader, actively developing rationale for pairing regulated cell death with checkpoint blockade across solid tumors^43^. The same metabolic theme extends to two further classifier genes, SLC39A4 (ZIP4) and DCLK1, both of which have documented, druggable roles in proliferation and stemness in other carcinomas^44,45^. Because both tumor groups share squamous histology, these differences are more plausibly attributable to genotype-specific viral biology than to cell-of-origin, implying that HPV16 and HPV18 expose different, potentially druggable molecular dependencies that are obscured by genotype-agnostic “high-risk HPV” framing.

Within each genotype, we further found that samples distributed along an early-to-late viral transcriptional continuum, with early-, mixed-, and late-dominant profiles rather than sharp discrete clusters, consistent with the differentiation-coupled nature of the productive HPV life cycle, in which early genes (E1, E2, E4, E5) predominate in undifferentiated basal keratinocytes and late genes (L1, L2) are normally restricted to terminally differentiated suprabasal cells^46^. Because HPV oncoproteins reinforce differentiation arrest in malignant cells^47^, mixed profiles may reflect incompletely dedifferentiated tumor subregions or non-canonical late-promoter activity uncoupled from true differentiation. Bulk RNA-seq cannot distinguish these possibilities, and single-cell or spatial transcriptomics will be needed to resolve whether mixed-profile cells co-localize with residual differentiation markers or disperse independently of tissue architecture. This interpretive caution extends to the early-and late-gene transcripts underlying the score itself: genome integration is well established to disrupt E2 and derepress E6/E7, yet purely integrated genomes were a minority configuration in at least one physical-status profiling study of HPV16-positive tumors^48^, with episomal fractions varying substantially across cohorts and the relationship between physical status and oncogene expression itself inconsistent across studies^49^. E2/E5 signal in our data can therefore arise from episomal copies coexisting with integrated ones rather than indicating an absence of integration, and detectable L1/L2 transcripts do not necessarily reflect progression toward virion assembly, since both carry translation-suppressing regulatory elements and L1 has a documented non-structural role in enhancing E2-dependent replication^50^. The viral transcriptional score is therefore best understood as a phenotypic readout of overall viral activity rather than a direct proxy for any single genomic configuration.

## 5 Conclusions

To our knowledge, this is the first study to systematically define HPV transcriptional programs directly from transcriptomic data on the basis of relative early-and late-gene expression, and to show that these programs couple to genotype-specific host responses, indicating that HPV-positive tumors occupy a richer space of biological states than genotype and infection status alone. These findings show that HPV16 and HPV18 do not converge on a single, genotype-agnostic route to cervical carcinogenesis, but instead rewire host biology through partly distinct regulatory, metabolic, and cell-death circuitry, from AhR/polarity signaling and RTK–RAS–MAPK activity in HPV16 tumors to interferon-coupled cuproptosis susceptibility and inflammatory-metabolic reprogramming in HPV18 tumors, exposing candidate therapeutic vulnerabilities that are obscured when the two genotypes are treated as interchangeable. The additional observation that viral transcriptional state itself couples to distinct host processes within each genotype further indicates that HPV-positive tumors occupy a richer space of biological states than genotype alone, with potential relevance to patient stratification and treatment selection. More broadly, this work demonstrates the value of recovering viral transcriptional signals directly from standard bulk RNA-seq data, an approach that could extend naturally to other high-risk HPV genotypes and other virus-associated malignancies where strain-specific tumor biology remains obscured by aggregate analysis.

## CRediT authorship contribution statement

Marcela Akemi Ishihara: Writing – review & editing, Writing – original draft, Investigation, Formal analysis, Data curation, Conceptualization. Maiara M. Moro Valeriano: Writing – review & editing, Writing – original draft, Investigation, Formal analysis, Data curation, Conceptualization.Noilson Oliveira: Writing – review & editing, Formal analysis. Izabela Mamede: Writing – review & editing, Writing – original draft, Formal analysis. Lívia Cavalcanti de Morais Gama: Writing – review & editing, Writing – original draft, Formal analysis. André Ferreira Cunha: Writing – review & editing, Formal analysis. Ícaro Maia Santos de Castro: Writing – review & editing, Methodology. Francisco Vásquez: Writing – original draft. Milton Yutaka Nishiyama-Jr: Writing – original draft, Resources. João Carlos Setubal: Writing – review & editing. Eurico de Arruda Neto: Writing – review & editing. Roger Chammas: Writing – review & editing. Helder I. Nakaya: Supervision, Conceptualization, Writing – review & editing, Writing – original draft.

## Acknowledgments

We acknowledge the computational infrastructure provided by the Nucleus of Bioinformatics and Computational Biology at the Butantan Institute (FAPESP grant no. 2022/11601-4) and by the Computational Systems Biology Laboratory at INOVA-USP, where the bioinformatics analyses were performed. We also thank the respective administration teams for their support. We are particularly thankful to the following grants for fellows: MAI (FAPESP process 2026/06696-7), MMMV (CNPq 131779/2023-0), IMSC (FAPESP 20/05284-0 and 23/03951-8) and HIN (2018/14933-2 and 2023/18337-3).

## Supporting Information

The Meta-HPV pipeline is publicly available at https://github.com/cell-ai/MetaHPV. Raw RNA-seq data for the Curty et al. (2021) and Ruiz et al. (2018) cohorts are deposited at NCBI SRA under BioProject accessions PRJNA356774 and PRJNA636799, respectively.

---

**Figure S1.**
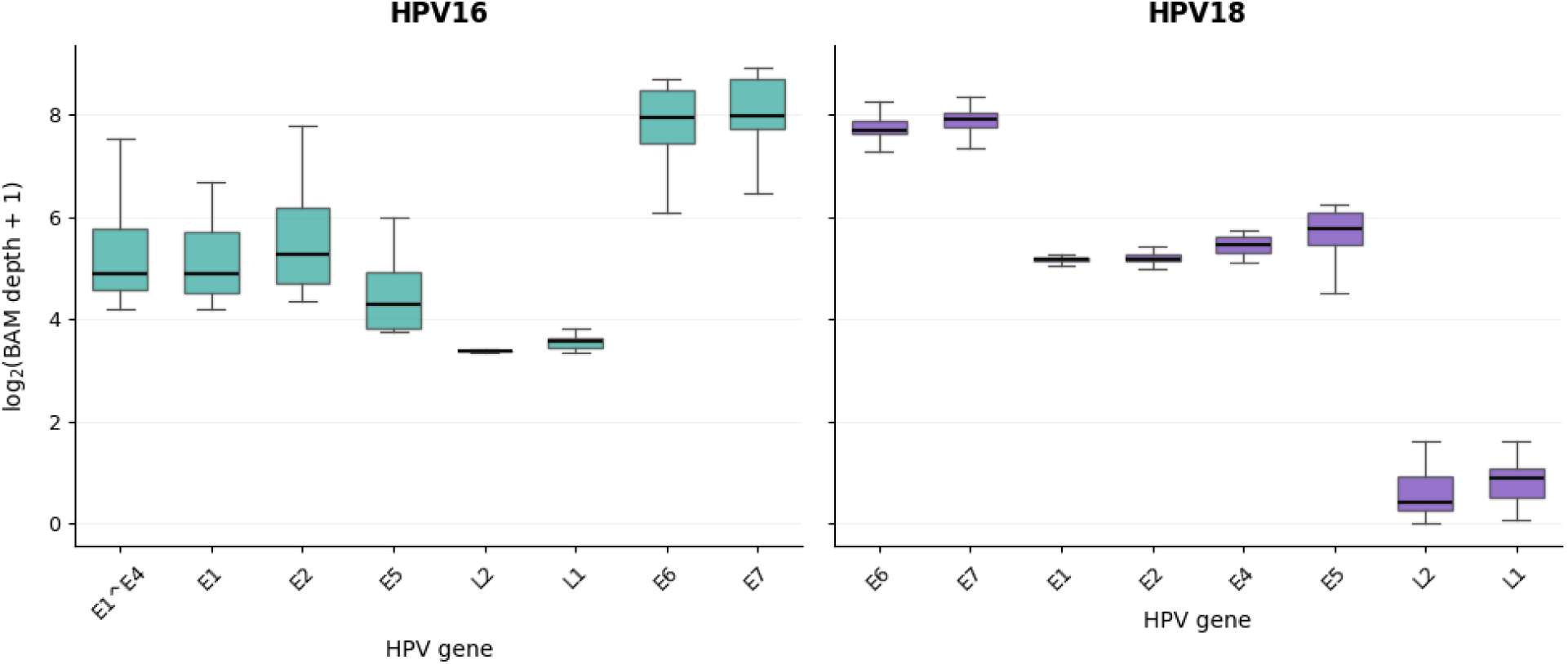
BAM sequencing depth across HPV genes. Boxplots show the per-base BAM sequencing depth across individual HPV genes in the new dataset, based on pooled PE and SE alignments. Depth is shown as log□(BAM depth + 1). The center line indicates the median, boxes represent the interquartile range, and whiskers extend to 1.5 × IQR; outliers are not shown. Teal and purple indicate HPV16 and HPV18, respectively.

## References

1. Walboomers, J. M. M. et al. Human papillomavirus is a necessary cause of invasive cervical cancer worldwide. J. Pathol. 189, 12–19 (1999).

2. Wu, J. et al. Global burden of cervical cancer: current estimates, temporal trend and future projections based on the GLOBOCAN 2022. J. Natl. Cancer Cent. 5, 322–329 (2025).

3. McBride, A. A. Human papillomaviruses: diversity, infection and host interactions. Nat. Rev. Microbiol. 20, 95–108 (2022).

4. Wei, F., Georges, D., Man, I., Baussano, I. & Clifford, G. M. Causal attribution of human papillomavirus genotypes to invasive cervical cancer worldwide: a systematic analysis of the global literature. The Lancet 404, 435–444 (2024).

5. Wang, Q., Song, R., Zhao, C., Liu, H. & Et., A. HPV16 E6 promotes cervical cancer cell migration and invasion by downregulation of NHERF1. Int. J. Cancer (2018) doi:10.1002/ijc.31876.

6. Wasson, C. W. et al. Human papillomavirus type 18 E5 oncogene supports cell cycle progression and impairs epithelial differentiation by modulating growth factor receptor signalling during the virus life cycle. Oncotarget 8, 103581–103600 (2017).

7. Reynders, C. et al. Prevalence and genotype distribution of human papillomavirus in cervical adenocarcinoma (usual type and variants): A systematic review and meta-analysis. J. Med. Virol. 95, e29190 (2023).

8. Ebisch, R. M. F. et al. Screening for persistent high-risk HPV infections may be a valuable screening method for young women; A retrospective cohort study. PLOS ONE 13, e0206219 (2018).

9. Im, S. S., Wilczynski, S. P., Burger, R. A. & Monk, B. J. Early Stage Cervical Cancers Containing Human Papillomavirus Type 18 DNA Have More Nodal Metastasis and Deeper Stromal Invasion. Clin. Cancer Res. 9, 4145–4150 (2003).

10. Ure, A., Mukhedkar, D. & Arroyo Mühr, L. S. Using HPV-meta for human papillomavirus RNA quality detection. Sci. Rep. 12, 13058 (2022).

11. Mjelle, R., Castro, Í. & Aass, K. R. The viral landscape in metastatic solid cancers. Heliyon 11, e42548 (2025).

12. Wood, D. E., Lu, J. & Langmead, B. Improved metagenomic analysis with Kraken 2. Genome Biol. 20, 257 (2019).

13. Kim, D., Paggi, J. M., Park, C., Bennett, C. & Salzberg, S. L. Graph-based genome alignment and genotyping with HISAT2 and HISAT-genotype. Nat. Biotechnol. 37, 907–915 (2019).

14. Li, H. et al. The Sequence Alignment/Map format and SAMtools. Bioinformatics 25, 2078–2079 (2009).

15. Wood, D. E., Lu, J. & Langmead, B. Improved metagenomic analysis with Kraken 2. Genome Biol. 20, 257 (2019).

16. Gihawi, A. et al. Major data analysis errors invalidate cancer microbiome findings. mBio 14, e01607-23 (2023).

17. Liao, Y., Smyth, G. K. & Shi, W. featureCounts: an efficient general purpose program for assigning sequence reads to genomic features. Bioinformatics 30, 923–930 (2014).

18. Love, M. I., Huber, W. & Anders, S. Moderated estimation of fold change and dispersion for RNA-seq data with DESeq2. Genome Biol. 15, 550 (2014).

19. Curty, G. et al. Expression of Retroelements in Cervical Cancer and Their Interplay with HPV Infection and Host Gene Expression. Cancers 13, 3513 (2021).

20. Ruiz, F. J., et al. HPV transcript expression affects cervical cancer response to chemoradiation. JCI Insight 6, e138734 (2018).

21. Chen, S., Zhou, Y., Chen, Y. & Gu, J. fastp: an ultra-fast all-in-one FASTQ preprocessor. Bioinformatics 34, i884–i890 (2018).

22. Rohart, F., Eslami, A., Matigian, N., Bougeard, S. & Lê Cao, K.-A. MINT: a multivariate integrative method to identify reproducible molecular signatures across independent experiments and platforms. BMC Bioinformatics 18, 128 (2017).

23. Robinson, M. D., McCarthy, D. J. & Smyth, G. K. edgeR: a Bioconductor package for differential expression analysis of digital gene expression data. Bioinformatics 26, 139–140 (2010).

24. Garcia-Alonso, L., Holland, C. H., Ibrahim, M. M., Turei, D. & Saez-Rodriguez, J. Benchmark and integration of resources for the estimation of human transcription factor activities. Genome Res. 29, 1363–1375 (2019).

25. Garcia-Alonso, L., Holland, C. H., Ibrahim, M. M., Turei, D. & Saez-Rodriguez, J. Benchmark and integration of resources for the estimation of human transcription factor activities. Genome Res. 29, 1363–1375 (2019).

26. Liberzon, A. et al. The Molecular Signatures Database Hallmark Gene Set Collection. Cell Syst. 1, 417–425 (2015).

27. Kim, D., Paggi, J. M., Park, C., Bennett, C. & Salzberg, S. L. Graph-based genome alignment and genotyping with HISAT2 and HISAT-genotype. Nat. Biotechnol. 37, 907–915 (2019).

28. Milacic, M. et al. The Reactome Pathway Knowledgebase 2024. Nucleic Acids Res. 52, D672–D678 (2024).

29. Ure, A. E., Lagheden, C. & Arroyo Mühr, L. S. Metatranscriptome analysis in human papillomavirus negative cervical cancers. Sci. Rep. 12, 15062 (2022).

30. Arroyo Mühr, L. S., et al. Deep sequencing detects human papillomavirus (HPV) in cervical cancers negative for HPV by PCR. Br. J. Cancer 123, 1790–1795 (2020).

31. Bulk, S. et al. Preferential risk of HPV16 for squamous cell carcinoma and of HPV18 for adenocarcinoma of the cervix compared to women with normal cytology in The Netherlands. Br. J. Cancer 94, 171–175 (2006).

32. Hellner, K., Mar, J., Fang, F., Quackenbush, J. & Münger, K. HPV16 E7 oncogene expression in normal human epithelial cells causes molecular changes indicative of an epithelial to mesenchymal transition. Virology 391, 57–63 (2009).

33. Waghmare, I. & Kango-Singh, M. Loss of Cell Adhesion Increases Tumorigenic Potential of Polarity Deficient Scribble Mutant Cells. PLOS ONE 11, e0158081 (2016).

34. Zheng, J. H. et al. Defective expression of polarity protein Par3 promotes cervical tumorigenesis and metastasis. Eur. J. Gynaecol. Oncol. 38, 199–206 (2017).

35. Wang, P., Laster, K., Jia, X., Dong, Z. & Liu, K. Targeting CRAF kinase in anti-cancer therapy: progress and opportunities. Mol. Cancer 22, 208 (2023).

36. Lagström, S. et al. HPV16 and HPV18 type-specific APOBEC3 and integration profiles in different diagnostic categories of cervical samples. Tumour Virus Res. 12, 200221 (2021).

37. Sullivan, L. M., Smolkin, M. E., Frierson, H. F. & Galgano, M. T. Comprehensive evaluation of CDX2 in invasive cervical adenocarcinomas: immunopositivity in the absence of overt colorectal morphology. Am. J. Surg. Pathol. 32, 1608–1612 (2008).

38. Kim, E.-S. et al. Inflammatory lipid sphingosine-1-phosphate upregulates C-reactive protein via C/EBPβ and potentiates breast cancer progression. Oncogene 33, 3583–3593 (2014).

39. He, A., Wu, C., Xia, D. & Zhu, Y. CEBPB emerges as a key regulatory factor in human cancers through diverse molecular mechanisms and clinical implications. Discov. Oncol. 16, 2278 (2025).

40. Wang, L. et al. Site-specific HPV18 integration facilitates cervical carcinogenesis through metabolic reprogramming-induced dysfunction of the SpHK1/S1P/S1PR1 pathway. Cell Death Dis. 17, 24 (2026).

41. Mandal, R. et al. The non-apoptotic function of Caspase-8 in negatively regulating the CDK9-mediated Ser2 phosphorylation of RNA polymerase II in cervical cancer. Cell. Mol. Life Sci. 79, 597 (2022).

42. Lei, G. et al. Cuproptosis-immunity crosstalk informs strategy to overcome immunotherapy resistance. Cell (2026) doi:10.1016/j.cell.2026.05.036.

43. Li, J. et al. Immunogenic cuproptosis in cancer immunotherapy via an in situ cuproptosis-inducing system. Biomaterials 319, 123201 (2025).

44. Li, M. et al. Aberrant expression of zinc transporter ZIP4 (SLC39A4) significantly contributes to human pancreatic cancer pathogenesis and progression. Proc. Natl. Acad. Sci. 104, 18636–18641 (2007).

45. Chandrakesan, P. et al. Regulatory Roles of Dclk1 in Epithelial Mesenchymal Transition and Cancer Stem Cells. J. Carcinog. Mutagen. 7, 257 (2016).

46. Graham, S. V. Keratinocyte Differentiation-Dependent Human Papillomavirus Gene Regulation. Viruses 9, 245 (2017).

47. Talora, C. et al. Constitutively active Notch1 induces growth arrest of HPV-positive cervical cancer cells via separate signaling pathways. Exp. Cell Res. 305, 343–354 (2005).

48. Hong, D. et al. Viral E6 is overexpressed via high viral load in invasive cervical cancer with episomal HPV16. BMC Cancer 17, 136 (2017).

49. Vinokurova, S. et al. Type-Dependent Integration Frequency of Human Papillomavirus Genomes in Cervical Lesions. Cancer Res. 68, 307–313 (2008).

50. Siddiqa, A. et al. The human papillomavirus type 16 L1 protein directly interacts with E2 and enhances E2-dependent replication and transcription activation. J. Gen. Virol. 96, 2274– 2285 (2015).

